# Defining metabolic flexibility in hair follicle stem cell induced squamous cell carcinoma

**DOI:** 10.1101/2023.10.16.562128

**Authors:** C Galvan, A Flores, V Cerrillos, I Avila, C Murphy, W Zheng, TT To, HR Christofk, WE Lowry

**Affiliations:** Department of Molecular Cell and Developmental Biology, UCLA; Department of Biological Chemistry, DGSOM, UCLA; Department of Medicine, DGSOM, UCLA; Molecular Biology Institute, UCLA; Broad Stem Cell Research Center, UCLA; Jonsson Comprehensive Cancer Center, UCLA

## Abstract

Among the numerous changes associated with the transformation to cancer, cellular metabolism is one of the first discovered and most prominent[1, 2]. However, despite the knowledge that nearly every cancer is associated with the strong upregulation of various metabolic pathways, there has yet to be much clinical progress on the treatment of cancer by targeting a single metabolic enzyme directly[3–6]. We previously showed that inhibition of glycolysis through lactate dehydrogenase (LDHA) deletion in cancer cells of origin had no effect on the initiation or progression of cutaneous squamous cell carcinoma[7], suggesting that these cancers are metabolically flexible enough to produce the necessary metabolites required for sustained growth in the absence of glycolysis. Here we focused on glutaminolysis, another metabolic pathway frequently implicated as important for tumorigenesis in correlative studies. We genetically blocked glutaminolysis through glutaminase (GLS) deletion in cancer cells of origin, and found that this had little effect on tumorigenesis, similar to what we previously showed for blocking glycolysis. Tumors with genetic deletion of glutaminolysis instead upregulated lactate consumption and utilization for the TCA cycle, providing further evidence of metabolic flexibility. We also found that the metabolic flexibility observed upon inhibition of glycolysis or glutaminolysis is due to post-transcriptional changes in the levels of plasma membrane lactate and glutamine transporters. To define the limits of metabolic flexibility in cancer initiating hair follicle stem cells, we genetically blocked both glycolysis and glutaminolysis simultaneously and found that frank carcinoma was not compatible with abrogation of both of these carbon utilization pathways. These data point towards metabolic flexibility mediated by regulation of nutrient consumption, and suggest that treatment of cancer through metabolic manipulation will require multiple interventions on distinct pathways.

## Introduction

Cutaneous Squamous Cell Carcinoma (SCC) is known to initiate in the epidermis due to an accumulation of mutations in genes such as Ras, Notch, P53, etc[8–11]. We and others showed that these cancers can arise due to transformation particularly of hair follicle stem cells (HFSCs) using murine transgenic and chemically induced carcinogenesis protocols[12, 13]. In humans these cancers are typically treated through surgical resection, but if left untreated can metastasize and be lethal. SCC can also arise in other areas that are much more difficult to treat such as in Head and Neck SCC, leading to much higher rates of mortality[9, 14–16]. Glutaminolysis has emerged as a key metabolic pathway in a variety of cancer models[17, 18]. Glutamine is imported into cells via plasma membrane glutamine transporters and can be converted to glutamate through the action of glutaminase (GLS) enzymes in the cytoplasm or the mitochondria[19]. Glutamate can be converted to alpha-ketoglutarate which can enter into the TCA cycle to power oxidative phosphorylation and production of copious ATP in the mitochondria. In addition, recent data show that many human tumors consume lactate through monocarboxylate transporters, convert lactate into pyruvate by lactate dehydrogenase, and utilize the lactate-generated pyruvate anapleurotically in the TCA cycle. Therefore, cancer cells can either put carbons into the TCA cycle either through uptake of glucose, lactate or glutamine, all of which have been shown to be upregulated in many human cancers. Despite numerous studies suggesting that glutaminolysis could be a driver of tumorigenesis, this has yet to be tested genetically *in vivo* in murine cancer models, particularly in SCC. To date, efforts to block glutaminolysis with small molecule inhibition of GLS activity have not yet led to clinically available therapies for patients despite significant effort[18, 20–22].

SCC and essentially every other solid tumor are known to show evidence of a metabolic transition known as the Warburg effect where cancer cells choose to increase glucose uptake as a nutrient and use it to produce lactate leading to acidification of tissue, a process also known as aerobic glycolysis[1, 2]. This observation has led to significant effort to block various targets in the glucose utilization pathway for the treatment of cancer. However, to date, these approaches have not been successful[5]. We previously showed that genetic deletion of Lactate Dehydrogenase A (LDHA) in cancer initiating cells of the epidermis led to a dramatic decrease in the Warburg Effect as measured by glucose uptake and lactate production[7]. However, SCC formed despite lack of LDHA, suggesting that cancer cells do not necessarily rely on glucose metabolism for their growth and transformation. Instead, these data raised the possibility that cancer cells show metabolic flexibility which allows them to grow by upregulating alternative pathways that generate ATP and biosynthesis of key materials to allow for increased proliferation. As potential evidence of flexibility, we also showed that tumors lacking LDHA activity exhibited increased glutamine consumption and glutaminolysis[7], however we did not experimentally investigate whether this increased glutamine metabolism enabled tumor growth in the absence of LDHA.

In the current study, we use a well-established murine model of SCC coupled with genetic deletion of LDHA and GLS to test the limits of cancer metabolic flexibility[23, 24]. In this model, genetic manipulation is inducible in the HFSCs, which are known to initiate SCC in murine epidermis[12, 25]. Coupled with chemical carcinogenesis, this model allows for the deletion of metabolic activity just prior to induction of oncogenesis in adult mice. Because the tumorigenesis begins at the skin surface, the entire process is tractable over time allowing for detailed quantification of oncogenesis and precise measurement of the role various metabolic pathways play in this process. We exploited this model to probe the role of glutaminolysis in SCC initiation or progression, and in doing so define metabolic flexibility in SCC as well as the limits of that flexibility.

## Results

Glutamine metabolism (Fig 1A) has previously been implicated as a key metabolic activity in tumor progression in a variety of cancer models[17, 21, 22, 26–28]. Here we sought to understand whether glutamine metabolism plays a role in tumor initiation or progression of SCC. We previously demonstrated that HFSCs serve as cells of origin for SCC, and that the tumors formed share many physiological and metabolic similarities with SCC formed in human skin[12, 29, 30]. First, we performed LCMS-based metabolomics to measure the relative levels of metabolites in HFSC-induced SCC and found elevated levels of several metabolites involved in glutamine metabolism (Fig 1B). Using our own transcriptional data we found many of the genes involved in glutaminolysis and glutamine metabolism were elevated in murine SCC derived from HFSCs (Fig 1C). We then examined cancer genome data (GENT2) to assess the relative mRNA levels of glutaminase, the enzyme that converts glutamine to glutamate as the first step of glutaminolysis, in normal versus tumorigenic tissues, which pointed towards higher GLS expression in many tumor types (Fig 1D). In skin tumors, GLS was also upregulated along with the glutamine transporter ASCT2, and GOT2 and ASNS, two other key enzymes in glutamine metabolism (Fig 1E). In a model of HFSC-induced tumorigenesis, we found that deletion of LDHA did not dramatically affect tumor production, but did show evidence of metabolic compensation by glutaminolysis (Fig 1F-H). On the other hand, transcriptional analysis of LDHA-null tumors did not show changes in expression of genes related to glutaminolysis raising the question of how this compensation was mediated (Fig S1).

**FIGURE 1.**
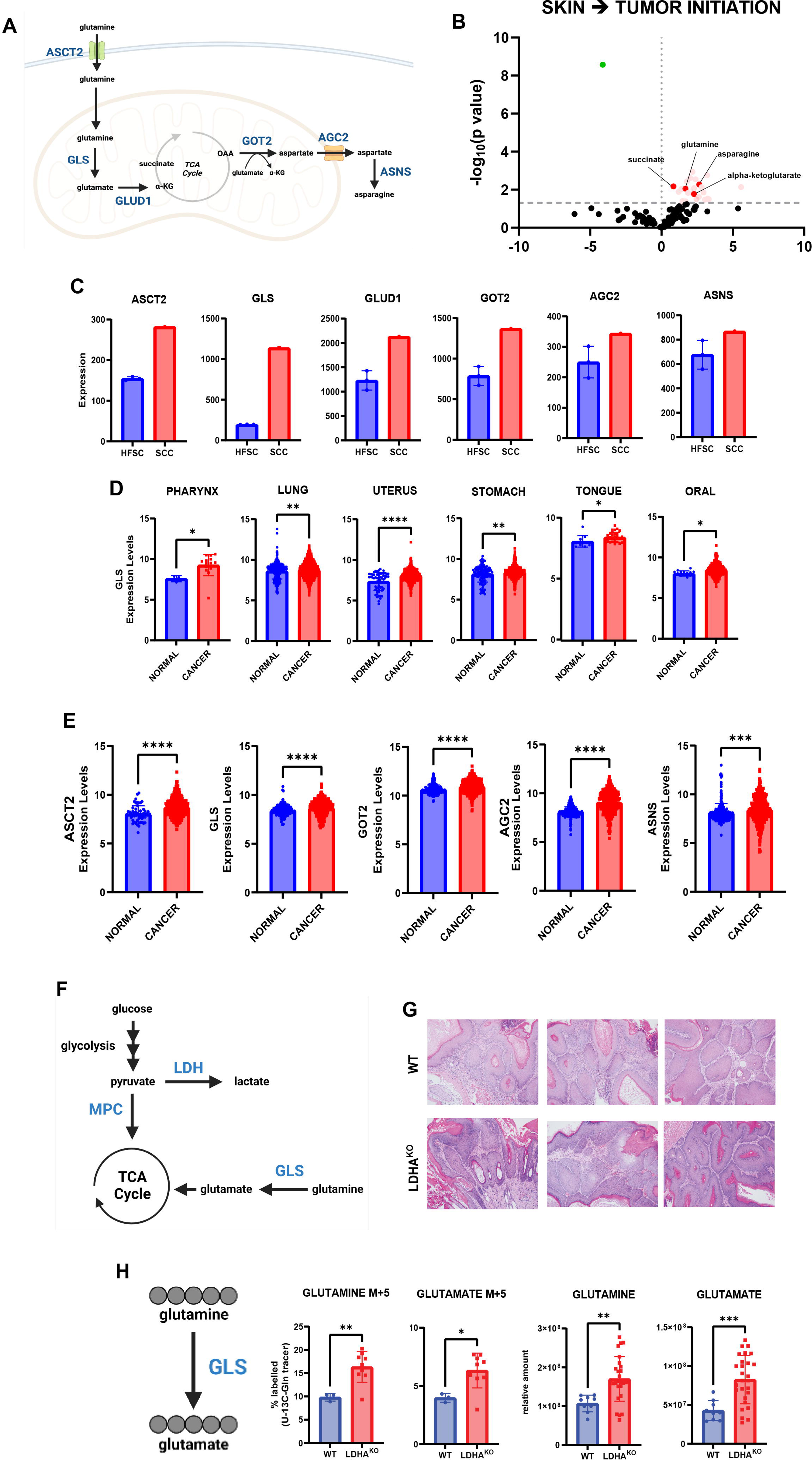
a) Schematic showing glutamine metabolism, glutamine being metabolized into TCA cycle and asparagine synthesis from TCA-cycle-derived aspartate. b) Metabolomic pool volcano plot of normal skin vs. papilloma (benign tumor) initiated by DMBA/TPA skin chemical carcinogenesis. Dashed lines indicate adjusted p≤0.05 or log_2_(fold change) ≥0 or ≤0. Colored dots represent metabolites significantly increasing (red) or decreasing (green) during the skin to tumor transition. c) ASCT2, GLS, GLUD1, GOT2, AGC2, and ASNS gene expression in HFSC and SCC. d) GLS gene expression in pharynx n = 3 (normal), n = 15 (cancer); lung n = 508 (normal), n = 2362 (cancer); uterus n = 58 (normal), n = 432 (cancer); stomach n = 117 (normal), n = 1028 (cancer); tongue n = 11 (normal), n = 30 (cancer); oral n = 15 (normal), n = 371 (cancer) based on the Gene Expression database of Normal and Tumor tissue 2 (GENT2) data analysis. Statistical significance (*p<0.05; **p<0.01; ***<0.001; ****p<0.0001) was calculated using a two-tailed t test. e) ASCT2, GLS, GOT2, AGC2, and ASNS gene expression in skin n = 263 (normal), n = 547 (cancer) based on GENT2 data analysis. Statistical significance (*p<0.05; **p<0.01; ***<0.001; ****p<0.0001) was calculated using a two-tailed t test. f) Schematic of enzymes used in glycolysis, lactate production, and glutaminolysis. g) Tumors from *WT* and *LDHA^KO^* mice stained for hematoxylin and eosin (H&E). h) Schematic of fully labeled glutamine isotopomer conversion. Data represents percent of M5 labeled glutamine and M5 labeled glutamate in tumors (n=3 (*WT)*, n= 9 (*LDHA^KO^*)) after ^13^C_5_-glutamine infusion. Statistical significance (*p<0.05; **p<0.01; ***<0.001; ****p<0.0001) was calculated using a two-tailed t test. Metabolic pool data representing relative amounts of glutamine and glutamate in tumors (n=9 (*WT)*, n= 27 (*LDHA^KO^*)). Statistical significance (*p<0.05; **p<0.01; ***<0.001; ****p<0.0001) was calculated using a two-tailed t test.

Since SCC tumors showed increased glutaminolysis-related gene expression, we sought to determine the role of glutaminolysis in the initiation or progression of SCC through deletion of GLS, the enzyme that converts glutamine to glutamate as the first step in glutaminolysis, in HFSCs prior to initiating tumorigenesis. Crossing mice floxed for GLS (GLS1 fl/fl, Jax Labs) with mice transgenic for K15CrePR allowed for an inducible deletion of GLS in HFSCs upon administration of the progesterone inhibitor Mifepristone (Fig 2A). To induce tumorigenesis, we relied on the established chemical carcinogenesis protocol using DMBA as a mutagen followed by repeated stimulation of proliferation by TPA[24]. Both *WT* and *GLS^KO^* models produced papilloma, well-differentiated SCC, moderately differentiated SCC, and keratoacanthoma (Fig 2B). We quantified time to tumor formation, number of tumors, and volume of tumors, but did not detect any significant difference in cancer formation in mice with or without HFSC GLS expression (Fig 2C). On the other hand, we did find that 6% of tumors that formed in mice with GLS deletion in cancer cells of origin became necrotic, which we did not observe in animals with GLS activity (Fig 2C).

**FIGURE 2.**
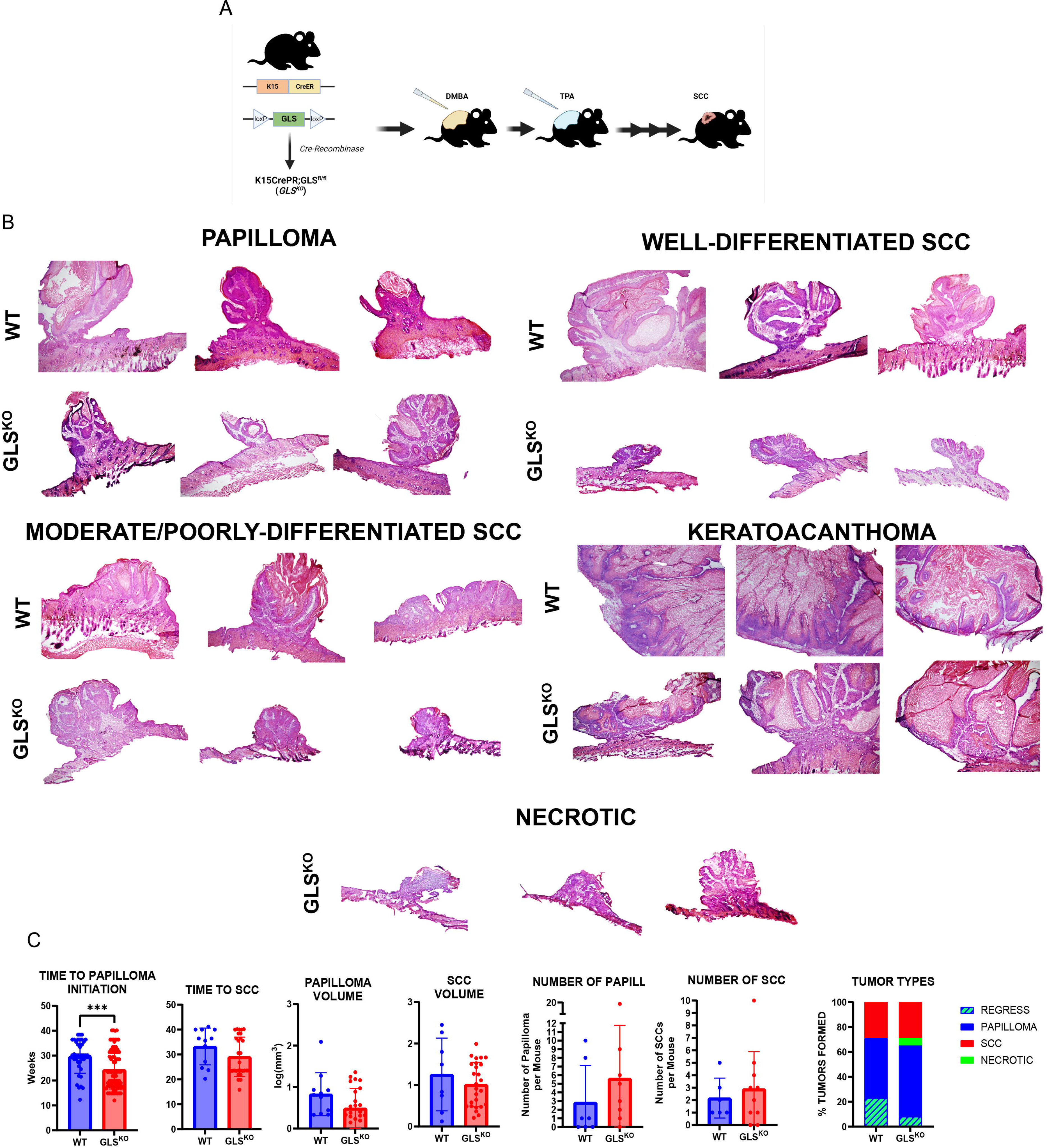
a) Schematic of transgenic mice used to knock out GLS in HFSCs coupled with topical SCC chemical carcinogenesis using DMBA and TPA. b) Dorsal tumors from *WT* and *GLS^KO^* mice stained for hematoxylin and eosin (H&E) at 4X magnification. c) Quantification of time to papilloma (n=40 (*WT)*, n=88 (*GLS^KO^*)) initiation and SCC (n=12 (*WT)*, n=29 (*GLS^KO^*)) formation. Each data point represents a tumor of that genotype. Quantification of volume of papilloma (n=11 (*WT)*, n=25 (*GLS^KO^*)) and SCC (n=8 (*WT)*, n=26 (*GLS^KO^*)). Each data point represents a tumor of that genotype. Quantification of the number of papilloma (n=7 (*WT)*, n=8 (*GLS^KO^*)) and SCC (n=6 (*WT)*, n=10 (*GLS^KO^*)) formed per mice. Each data point represents a mouse of that genotype. Data shown represents tumors present at the end of the experiment. Quantification of percent and types of tumors formed per genotype. *WT* (papilloma=48%; SCC=30%; regress=23%; necrotic=0%) and *GLS^KO^* (papilloma=57%; SCC=30%; regress=8%; necrotic=6%). Data shown represents tumor quantifications from the beginning to the end of the experiment.

To examine whether the genetic deletion of GLS in HFSCs effectively created tumors lacking GLS activity, we used immunostaining, GLS activity assays and LCMS-based metabolomics measurements. Immunostaining of *WT* tumors showed high GLS expression particularly on the epithelial edge of tumors formed after DMBA-TPA, whereas immunostaining of *GLS^KO^* tumors resulted in negligible GLS staining (Fig 3A). We also adapted a GLS activity assay to measure the relative activity of the enzyme in protein lysates generated from *WT* vs *GLS^KO^* tumors, and found a strong decrease in GLS activity in the *GLS^KO^* tumors (Fig 3A). While it is clear that our genetic strategy to eliminate GLS activity from tumors was successful, we did find cells that were strongly positive for GLS expression in the mesenchyme surrounding the nascent tumors (Fig S2A). These CD45+ cells were also positive for CD11b, suggesting they are macrophages (Fig S2B-C). Because our genetic strategy was not designed to target immune cells, it is not surprising to find GLS-positive cells within the mesenchyme, and these will be the subject of future investigations.

**FIGURE 3.**
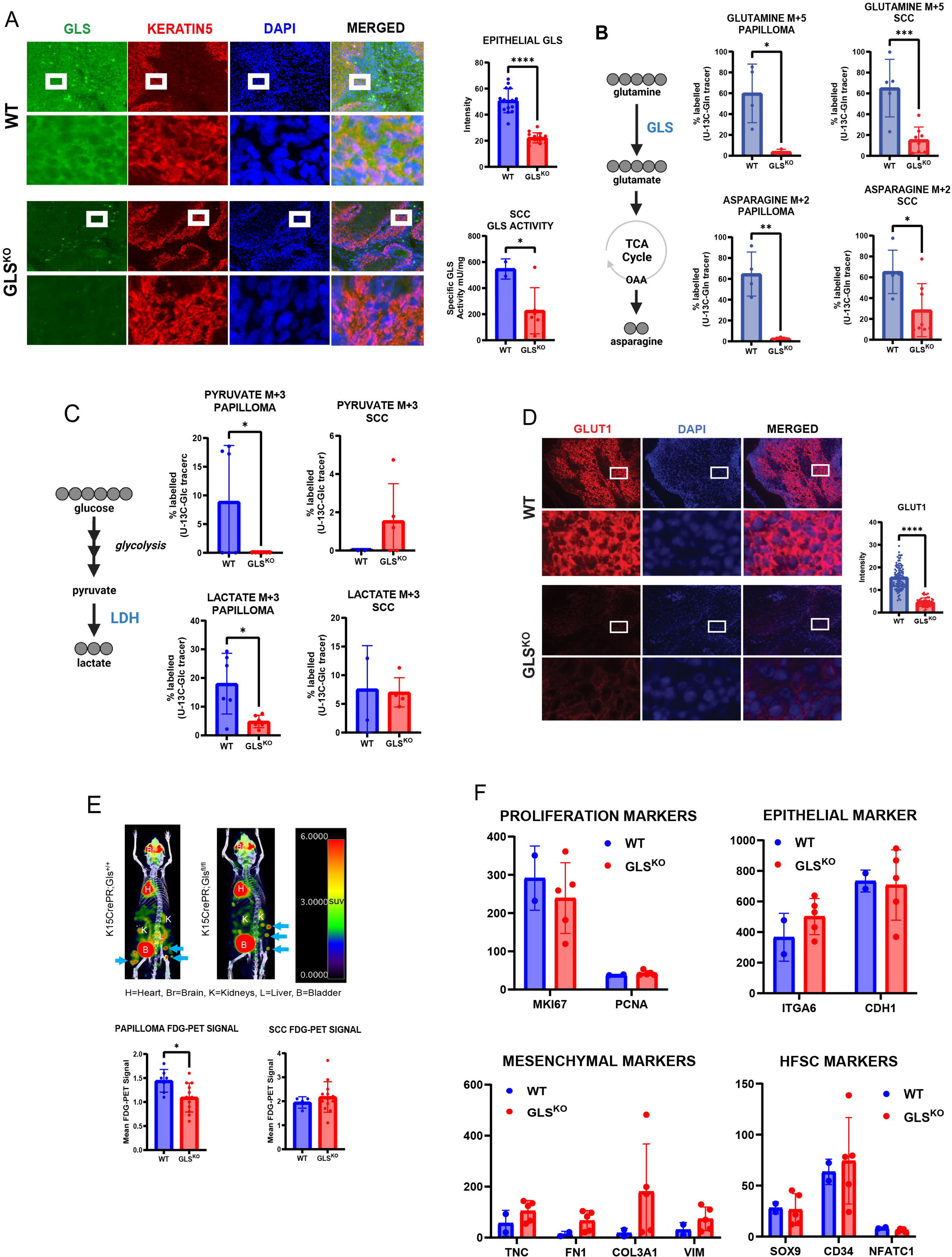
a) WT or *GLS^KO^* SCC immunostaining for GLS and KERATIN5, an epidermal marker. Cell nuclei were stained with DAPI. Quantification of mean intensity epithelial GLS fluorescence in *WT* (n=15) and *GLS^KO^* (n=15) SCCs. GLS activity in *WT* (n=2) and *GLS^KO^*(n=6) SCCs lysates. Statistical significance (*p<0.05; **p<0.01; ***<0.001; ****p<0.0001) was calculated using a two-tailed t test. b) Schematic of fully labeled glutamine isotopomer conversion. Data represents percent of M5 labeled glutamine and M2 labeled asparagine in papilloma (n=4 (*WT)*, n=3 (*GLS^KO^*)) and SCC (n=5 (*WT)*, n=8 (*GLS^KO^*)) after ^13^C_5_-glutamine infusion. c) Schematic of fully labeled glucose isotopomer conversion. Data represents percent of M3 labeled pyruvate and M3 labeled lactate in papilloma (n=6 (*WT)*, n=6 (*GLS^KO^*)) and SCC (n=2 (*WT)*, n=5 (*GLS^KO^*)) after ^13^C_6_-glucose infusion. d) *WT* or *GLS^KO^* SCC immunostaining for glucose transporter, GLUT1. Cell nuclei were stained with DAPI. Quantification of mean intensity GLUT1 fluorescence in *WT* (n=91) and *GLS^KO^* (n=56) SCCs. Statistical significance (*p<0.05; **p<0.01; ***<0.001; ****p<0.0001) was calculated using a two-tailed t test. e) Mean ^18^F-FDG SUV signal of papilloma (n=7 (*WT)*, n=12 (*GLS^KO^*)) and SCC (n=4 (*WT)*, n=15 (*GLS^KO^*)). H=heart; Br=brain; K=kidneys; L=liver; B=bladder. Statistical significance (*p<0.05; **p<0.01; ***<0.001; ****p<0.0001) was calculated using a two-tailed t test. f) RNA-seq data of *WT* (n=2) or *GLS^KO^* (n=5) tumors showing transcription levels of proliferation, epithelial, mesenchymal, and HFSC markers.

To investigate whether GLS-deleted tumors show evidence of metabolic changes despite relative lack of phenotypic change, we performed metabolic tracing with ^13^C-glutamine and LCMS-based metabolomics. Tumors with labeled glutamine showed that both the oxidative and reductive pathways for glutamine utilization were abrogated in GLS deleted tumors (Fig 3B), consistent with a loss of GLS activity. On the other hand, metabolomics also showed consistent decreases in glucose conversion to pyruvate and lactate in the absence of GLS in papilloma, but not in SCC (Fig 3C). Immunostaining for GLUT1, a glucose transporter known to be upregulated in SCC, showed diminished expression (Fig 3D). Furthermore, FDG-PET imaging, which measures the rate of FDG uptake as a proxy for glucose uptake in tumors of live animals, also showed signs of decreased glucose uptake in the absence of GLS specifically in papilloma (Fig 3E)[7]. RNA-seq of *WT* and *GLS^KO^* tumors showed a relatively small number of gene expression differences caused by loss of GLS activity. When looking particularly at genes related to proliferation, EMT, or stemness, there are no significant changes (Fig 3F).

Since ^13^C-glutamine and ^13^C-glucose tracing in *GLS^KO^* tumors revealed decreased glutamine and glucose consumption, we examined whether *GLS^KO^* tumors increased consumption of lactate, another abundant nutrient in circulation. Interestingly, ^13^C-lactate tracing revealed a large increase in lactate uptake in *GLS^KO^*tumors (Fig 4A). In addition, it appeared as though lactate was able to be processed both for gluconeogenesis and the TCA cycle, suggesting that increased lactate uptake was able to power the TCA cycle to compensate for the loss of glutaminolysis. We looked for evidence of changes in lactate transporter expression in *GLS^KO^* tumors by staining for MCT1 and MCT4. These two transporters are known to allow both lactate and pyruvate to traverse the plasma membrane in both directions as needed. We found that MCT1 expression was unchanged in *GLS^KO^* tumors, but MCT4 was upregulated in *GLS^KO^*tumors, providing an explanation for the increased lactate uptake and utilization in *GLS^KO^* tumors (Fig 4B-C). However, neither MCT1 nor MCT4 were differentially expressed at the RNA level, consistent with a post-transcriptional mechanism by which MCT4 protein is changed in these tumors (Fig 4D).

**FIGURE 4.**
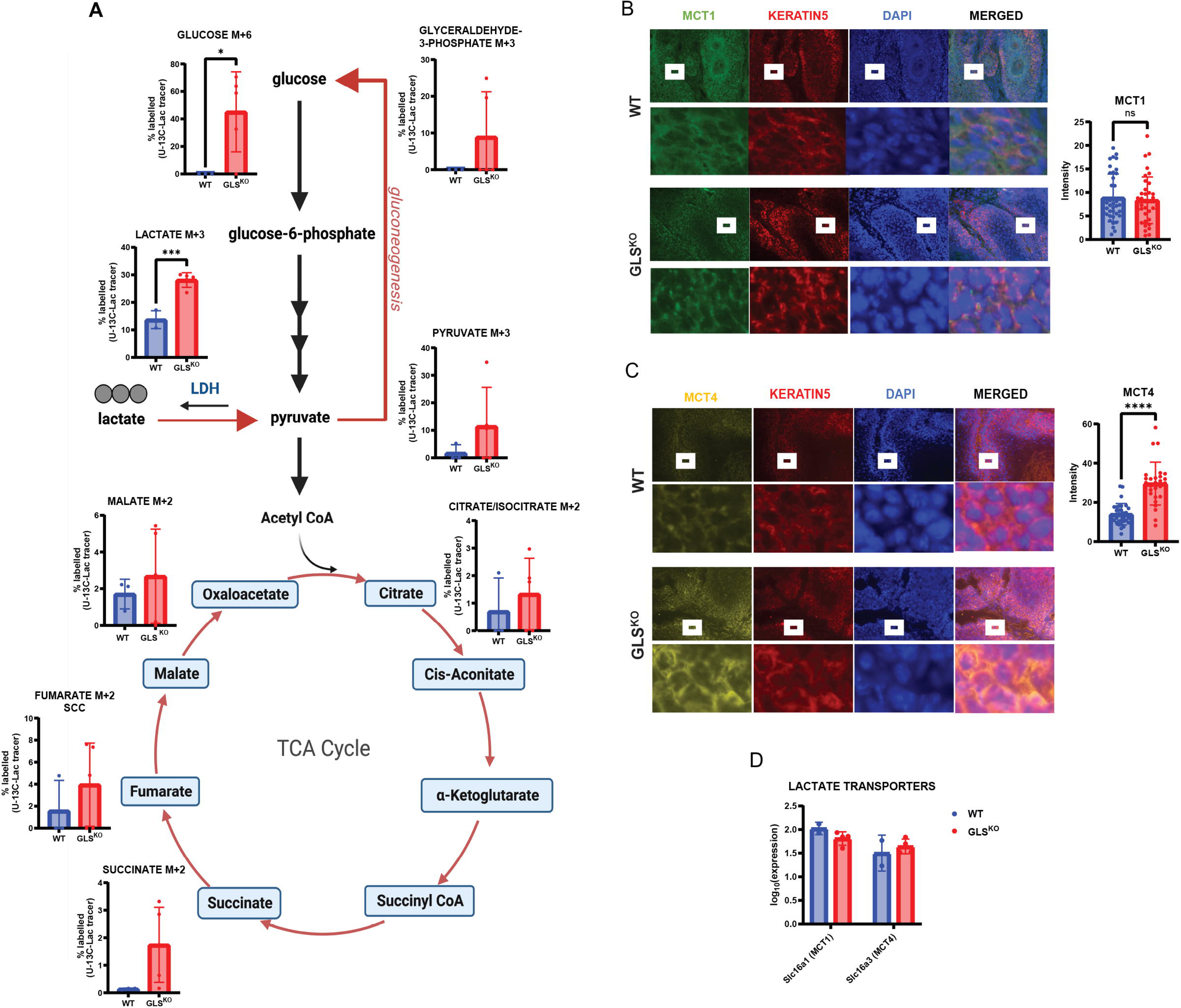
a) Schematic of fully labeled lactate isotopomer conversion. Data represents percent of M3 labeled lactate, M6 labeled glucose, M3 labeled glyceraldehyde-3-phosphate, M3 labeled pyruvate, M2 labeled citrate/isocitrate, M2 labeled succinate, M2 labeled fumarate, and M2 labeled malate in SCCs (n=3 (*WT)*, n=5 (*GLS^KO^*)) after ^13^C_3_-lactate infusion. b) *WT* or *GLS^KO^*SCC immunostaining for lactate transporter, MCT1. Cell nuclei were stained with DAPI. Quantification of mean intensity MCT1 fluorescence in *WT* (n=39) and *GLS^KO^* (n=36) SCCs. Statistical significance (*p<0.05; **p<0.01; ***<0.001; ****p<0.0001) was calculated using a two-tailed t test. c) *WT* or *GLS^KO^*SCC immunostaining for lactate transporter, MCT4. Cell nuclei were stained with DAPI. Quantification of mean intensity MCT4 fluorescence in *WT* (n=33) and *GLS^KO^* (n=27) SCCs. Statistical significance (*p<0.05; **p<0.01; ***<0.001; ****p<0.0001) was calculated using a two-tailed t test. d) RNA-seq data of *WT* (n=2) or *GLS^KO^* (n=5) tumors showing transcription levels of lactate transporters

These results suggesting metabolic flexibility in *GLS^KO^*tumors and capability to switch to a different carbon source through altering expression of nutrient transporters – particularly MCT4 - prompted us to examine metabolic flexibility and nutrient transporter expression in tumors initiated by HFSCs lacking LDHA. As described previously, *LDHA^KO^* tumors appeared pathologically identical to tumors expressing LDHA (Fig 1G). We pulsed mice bearing *LDHA^KO^*tumors with ^13^C-labeled glucose prior to tumor harvesting, and not surprisingly, found diminished tumor glucose uptake and conversion of glucose to lactate, and decreased tumor levels of glucose and lactate, confirming abrogation of glucose metabolism in the absence of LDHA activity described in our previous study (Fig 5A-B). In addition, we re-examined metabolomic data from tumors generated by HFSCs lacking the mitochondrial pyruvate carrier (MPC). In this model, tumorigenesis was also unaffected by blocking pyruvate oxidation, providing yet another example of metabolic flexibility. Increased glucose, lactate, glutamine and glutamate levels in MPC-null tumors suggested a potential upregulation of glycolysis and glutaminolysis (Fig 5C). These data, coupled with our observations about lactate transporter, MCT4, prompted us to ask whether glucose transporters are potentially dynamically regulated to mediate metabolic flexibility. We therefore immunostained for GLUT1, the glucose transporter, and found that GLUT1 protein expression at the cell membrane was strongly downregulated in *LDHA^KO^*tumors but upregulated in *MPC^KO^* tumors (Fig 5D and E).

**FIGURE 5.**
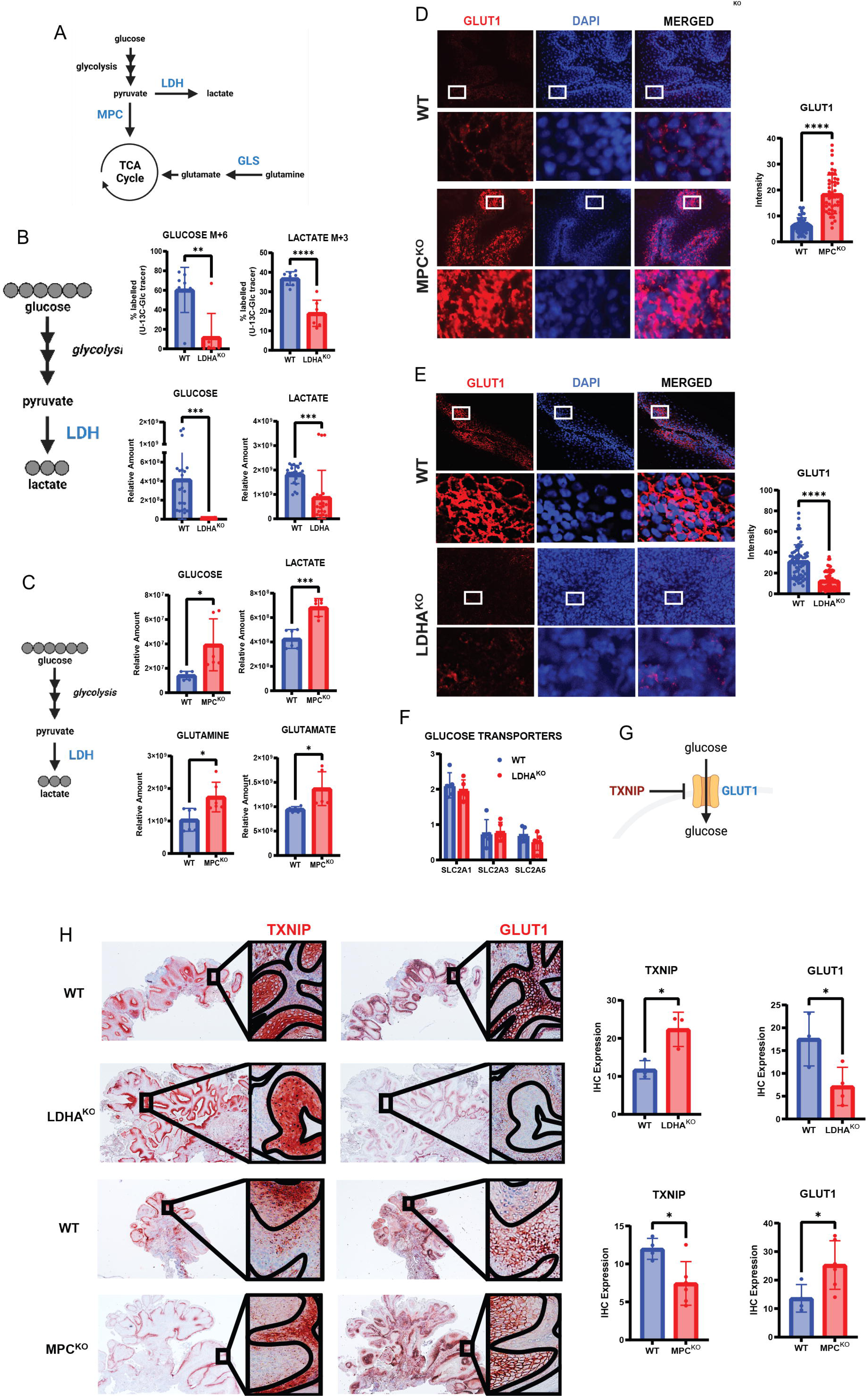
a) Schematic of glycolysis, lactate production, and glutaminolysis. b) Schematic of fully labeled glucose isotopomer conversion. Data represents percent of M6 labeled glucose and M3 labeled lactate in tumors (n=8 (*WT)*, n= 7 (*LDHA^KO^*)) after ^13^C_6_-glucose infusion. Metabolic pool data representing relative amounts of glucose and lactate in tumors (n=24 (*WT)*, n= 21 (*LDHA^KO^*)). Statistical significance (*p<0.05; **p<0.01; ***<0.001; ****p<0.0001) was calculated using a two-tailed t test. c) Metabolic pool data representing relative amounts of glucose, lactate, glutamine, and glutamate in tumors (n=6 (*WT)*, n=6 (*MPC^KO^*)). Statistical significance (*p<0.05; **p<0.01; ***<0.001; ****p<0.0001) was calculated using a two-tailed t test. d) *WT* or *MPC^KO^*SCC immunostaining for glucose transporter, GLUT1. Cell nuclei were stained with DAPI. Quantification of mean intensity GLUT1 fluorescence in *WT* (n=56) and *MPC^KO^* (n=49) SCCs. Statistical significance (*p<0.05; **p<0.01; ***<0.001; ****p<0.0001) was calculated using a two-tailed t test. e) *WT* or *LDHA^KO^*SCC immunostaining for glucose transporter, GLUT1. Cell nuclei were stained with DAPI. Quantification of mean intensity GLUT1 fluorescence in *WT* (n=61) and *LDHA^KO^* (n=65) SCCs. Statistical significance (*p<0.05; **p<0.01; ***<0.001; ****p<0.0001) was calculated using a two-tailed t test. f) RNA-seq data of *WT* (n=5) or *LDHA^KO^* (n=5) tumors showing transcription levels of glucose transporters g) Schematic of TXNIP inhibiting GLUT1 h) *WT*, *LDHA^KO^*or *MPC^KO^* tumor serial sections were probed for TXNIP and GLUT1. Lines in zoomed images mark boundaries of tissue structures and dashed lines highlight anti-coorelation between TXNIP and GLUT1. Quantification of TXNIP expression (n=3 (*WT)*, n=3 (*LDHA^KO^*)) and (n=4 (*WT)*, n=6 (*MPC^KO^*)) SCCs. Quantification of GLUT1 expression (n=3 (*WT)*, n=4 (*LDHA^KO^*)) and (n=4 (*WT)*, n=6 (*MPC^KO^*)) SCCs. Statistical significance (*p<0.05; **p<0.01; ***<0.001; ****p<0.0001) was calculated using a two-tailed t test.

Surprisingly, RNA-seq showed no changes in the expression of glucose transporter in LDHA-null tumors (Fig 5F) suggesting a post-transcriptional mechanism for the changes in GLUT1 protein levels. To identify such a mechanism, we looked at expression of TXNIP, which is known to regulate GLUT1 levels at the plasma membrane (Fig 5G). Therefore, we stained for these two proteins in the various tumor models described here to see if a correlation of expression of these proteins could serve to explain the membrane up- or downregulation of metabolite transporters in response to genetic manipulation of LDHA, or MPC. IHC for TXNIP and GLUT1 showed a remarkable anti-correlation as previously described, and quantification of total expression for both of these proteins showed that TXNIP is expressed much lower in MPC-null tumors, and the converse was true in LDHA-null tumors (Fig 5H).

We next immunostained for ASCT2, a plasma membrane glutamine transporter, in our models of tumors initiated by HFSCs. *GLS^KO^* tumors showed a significant decrease in ASCT2 protein further explaining the decrease in glutamine uptake and metabolism in these tumors (Fig 6A). ASCT2 protein at the membrane showed a strong increase in LDHA-null tumors, providing a potential mechanism by which glutaminolysis was induced in these tumors (Fig 6B). Profiling multiple *WT* and *GLS^KO^* tumors using RNA-seq analysis showed that ASCT2, nor other relevant transporters were differentially expressed (Fig 6C). ASCT2 is thought to form a complex with activated EGFR receptor [31], which is also known to be highly active in Ras driven SCC[32, 33]. We immunostained for active EGFR and indeed found strong expression at the cell membrane in SCC driven by DMBA/TPA (Fig 6D). In tumors generated by *GLS^KO^* HFSCs, phospho-EGFR expression appeared to be confined to the nucleus (Fig 6E). On the other hand, in tumors from the *LDHA^KO^* background, the phospho-EGFR was at the membrane and expressed at a higher level, similar to what was observed for ASCT2 (Fig 6F). Previous studies have shown that some tumors show nuclear EGFR expression [34], and this mechanism could explain why ASCT2 protein expression at the membrane disappears in the absence of GLS without any change in transcription (Fig 6G). Therefore, the elevated glutaminolysis observed in *LDHA^KO^* tumors, and diminished glutamine uptake observed in *GLS^KO^* tumors could be due to dynamic regulation of active EGFR at the membrane.

**FIGURE 6.**
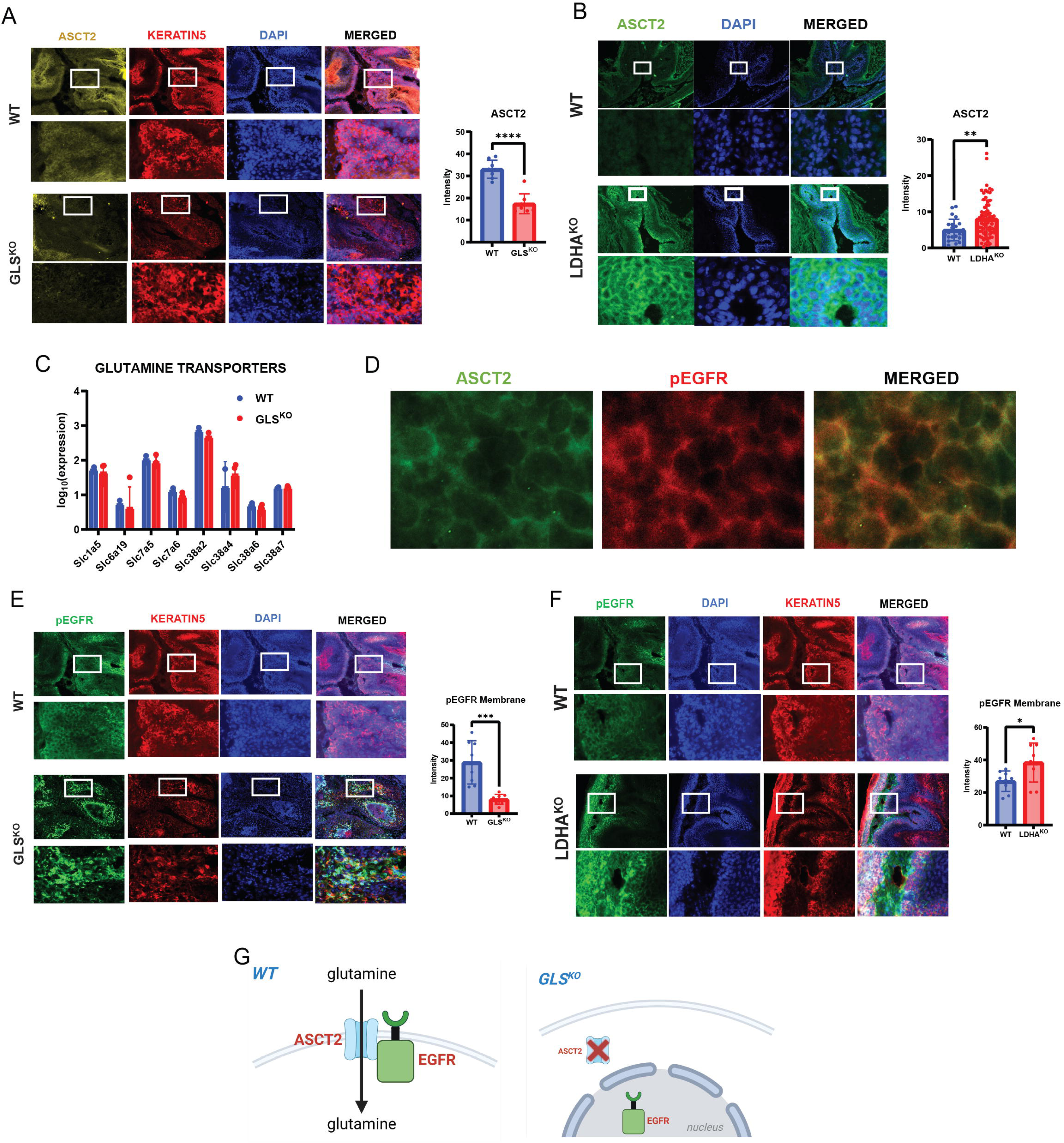
a) WT or *GLS^KO^* SCC immunostaining for glutamine transporter, ASCT2. Cell nuclei were stained with DAPI. Quantification of mean intensity ASCT2 fluorescence in *WT* (n=8) and *GLS^KO^* (n=8) SCCs. Statistical significance (*p<0.05; **p<0.01; ***<0.001; ****p<0.0001) was calculated using a two-tailed t test. b) *WT* or *LDHA^KO^*SCC immunostaining for glutamine transporter, ASCT2. Cell nuclei were stained with DAPI. Quantification of mean intensity ASCT2 fluorescence in *WT* (n=23) and *LDHA^KO^* (n=74) SCCs. Statistical significance (*p<0.05; **p<0.01; ***<0.001; ****p<0.0001) was calculated using a two-tailed t test. c) RNA-seq data of *WT* (n=2) or *GLS^KO^*(n=5) tumors showing transcription levels of glutamine transporters d) *WT* SCC immunostaining for ASCT2 and p-EGFR. e) *WT* or *GLS^KO^*SCC immunostaining for p-EGFR. Cell nuclei were stained with DAPI. Quantification of mean intensity p-EGFR fluorescence in *WT* (n=8) and *GLS^KO^* (n=8) SCCs. Statistical significance (*p<0.05; **p<0.01; ***<0.001; ****p<0.0001) was calculated using a two-tailed t test. f) *WT* or *LDHA^KO^*SCC immunostaining for p-EGFR. Cell nuclei were stained with DAPI. Quantification of mean intensity p-EGFR fluorescence in *WT* (n=9) and *LDHA^KO^*(n=9) SCCs. Statistical significance (*p<0.05; **p<0.01; ***<0.001; ****p<0.0001) was calculated using a two-tailed t test. g) Schematic proposing ASCT2 and EGFR localizations in *WT* and *GLS^KO^* SCCs.

The data from Figures 4-6 suggest that SCC-initiating cells have metabolic flexibility for carbon sources to power metabolic pathways, so we hypothesized that perhaps deletion of two carbon sources might be sufficient to starve cells attempting transformation. To test this hypothesis, we crossed animals floxed for both LDHA and GLS with K15CrePR transgenic mice in an attempt to abrogate both glucose utilization as well as glutaminolysis (Fig 7A). We then treated double floxed mice with DMBA/TPA in an attempt to induce tumorigenesis. After 20-30 weeks we routinely detected papilloma in *GLS^KO^LDHA^KO^*mice (Fig 7B), but never observed the formation of an SCC. By the end of the experiment, when the control animals had to be euthanized, there were no papilloma or SCC that truly lacked both LDHA activity and GLS expression in any of the *GLS^KO^LDHA^KO^*mice (Fig 7B). This suggests that those papilloma that lacked both LDHA and GLS that might have formed in *GLS^KO^LDHA^KO^* mice probably underwent regression. Careful chronological examination of tumorigenesis in the *WT* vs *GLS^KO^* vs *GLS^KO^LDHA^KO^*tumors showed that indeed, all tumors formed in *GLS^KO^LDHA^KO^*mice were benign papilloma which then either underwent necrosis or regression (Fig 7B-D). These data potentially define the limit of metabolic flexibility in SCC and point toward the utility of blocking multiple metabolic pathways to treat cancer.

**FIGURE 7.**
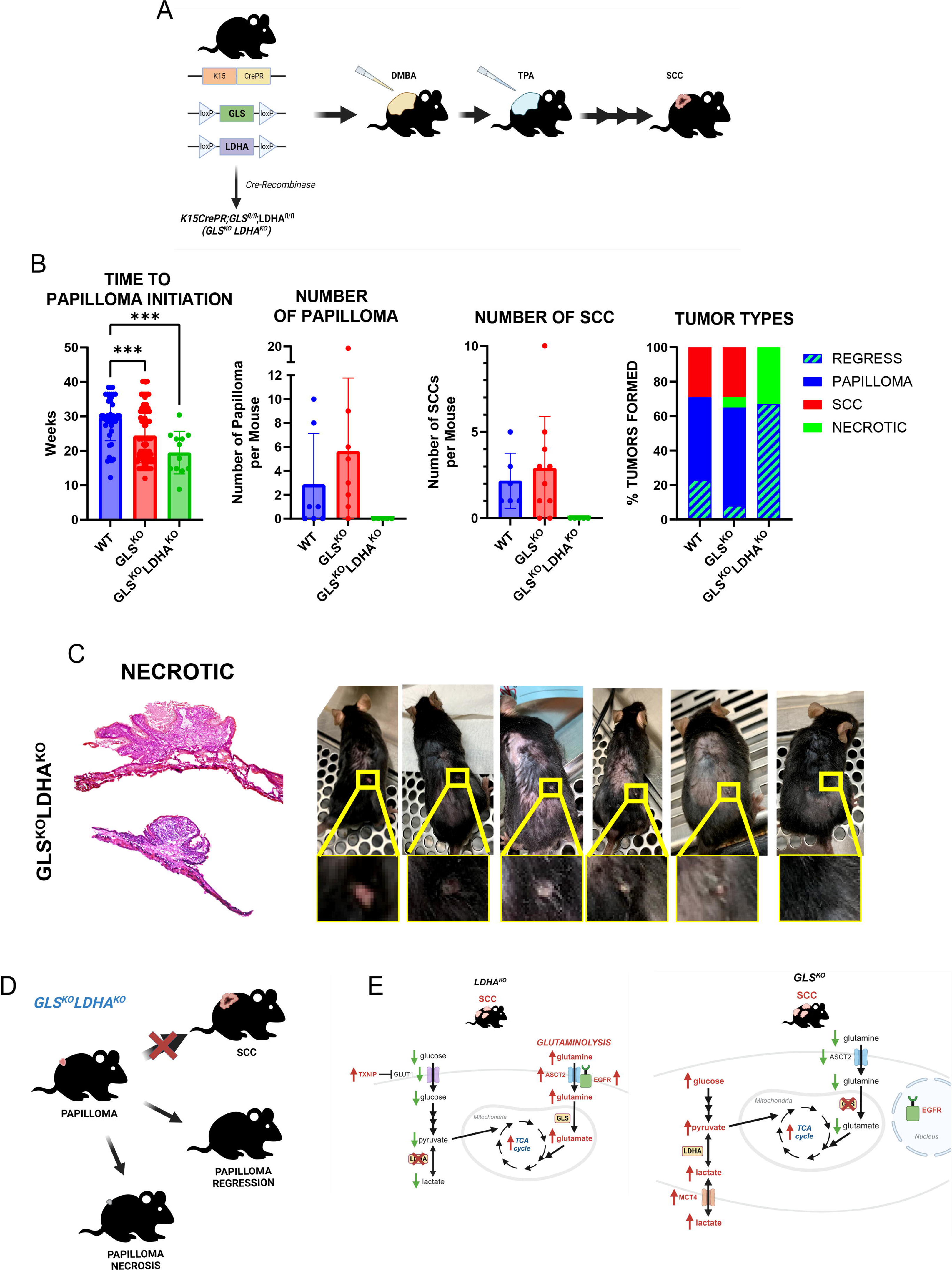
a) Schematic of transgenic mice used to knock out GLS and LDHA in HFSCs coupled with topical SCC chemical carcinogenesis using DMBA and TPA. b) Quantification of time to papilloma n=40 (*WT)*, n=88 (*GLS^KO^*), n=12 (*GLS^KO^LDHA^KO^*) initiation. Each data point represents a tumor of that genotype. Quantification of the number of papilloma (n=7 (*WT)*, n=8 (*GLS^KO^*), n=5 (*GLS^KO^LDHA^KO^*)) and SCC (n=6 (*WT)*, n=10 (*GLS^KO^*), n=5 (*GLS^KO^LDHA^KO^*)). Each data point represents a mouse of that genotype. Data shown represents tumors present at the end of the experiment. Quantification of percent and types of tumors formed per genotype. *WT* (papilloma=48%; SCC=30%; regress=23%; necrotic=0%), *GLS^KO^* (papilloma=57%; SCC=30%; regress=8%; necrotic=6%), and *GLS^KO^LDHA^KO^* (papilloma=0%; SCC=0%; regress=67%; necrotic=33%). c) Necrotic tumors from *GLS^KO^LDHA^KO^* mice stained for hematoxylin and eosin (H&E). Images of *GLS^KO^LDHA^KO^* mouse over time undergoing papilloma regression. d) Schematic of summary of phenotypic results for *GLS^KO^LDHA^KO^* mice. e) Schematic of proposed mechanisms of metabolic flexibility in *LDHA^KO^* and *GLS^KO^* HFSC-induced SCCs.

## Discussion

Together these data define the limits of metabolic flexibility of tumor initiating cells, and potentially point towards novel therapeutic combinatorial strategies to fight cancer initiation and progression. While previous studies argued that blockade of individual metabolic nodes could be an effective treatment strategy, our data argue otherwise. It is worth speculating that the difference between these outcomes could be due to the cancer models being employed, namely *in vitro* versus *in vivo*. Our data are consistent with studies showing that instead, multiple metabolic pathways must be targeted to overcome metabolic flexibility of cancer cells, including a previous *in vivo* study in lung cancer showing that tumors that are insensitive to inhibition of glycolysis or GLS alone, are in fact sensitive to the combination of both glycolysis and GLS inhibition [35–37].

*In vivo*, cancer cells potentially have access to a much more sophisticated environment due to the presence of vasculature, circulation, lymphatics, the nervous system, the immune system, etc. Therefore, it is possible that blockade of an individual pathway *in vitro* blocks cell growth simply because nutrients that allow for flexibility are not present in cell culture systems. As a result, therapeutic abrogation of cancer progression will probably require pharmacological targeting of more than one pathway as our genetic data presented here suggest. Fortunately, significant effort has been devoted to creating small molecule inhibitors of various metabolite uptake and utilization pathways such as CB839 (GLS), GSK2837808A (LDHA), UK5099 (MPC), AZD0095 (MCT4). All of these compounds show good safety profiles, however, none of these compounds by themselves effectively block tumor progression, but perhaps could be more efficacious in combination. It is worth noting that another approach with the drug DON is beginning to show promise in clinical trials[18, 38, 39]. However, this compound appears to target many proteins in the glutamine utilization pathway, and perhaps has targets outside the glutamine pathway, in which case would be consistent with the hypothesis that it is necessary to target multiple metabolic nodes to treat cancer.

Perhaps the most intriguing result from these experiments is the potential for metabolic flexibility to be mediated by regulation of transporters. Our data show that when LDHA activity is blocked, an increase of glutamine uptake coincides with upregulation of the ASCT2 transporter (Fig 7E). Conversely, when GLS activity was genetically blocked, lactate uptake was increased along with expression of MCT4 transporter at the cell membrane (Fig 7E). Furthermore, we expanded previous findings to demonstrate that the GLUT1 transporter is key to the promotion and diminution of glycolytic activity observed in LDHA and MPC deleted tumors. The regulation of these transporters did not appear to be at the transcriptional level, as RNA-seq failed to identify changes in RNA expression of any of these transporters. Our data suggest that interactions between these transporters and proteins such as EGFR and TXNIP mediate the regulation of cell membrane localization of these transporters, which then appear to drive metabolic flexibility. The mechanism proposed here demonstrates that cancer cells have a rather elegant means with which to compensate for loss of function of a metabolic pathway by simply putting more transporter at the membrane for an alternate pathway.

As a result, it is worth speculating that inhibition of nutrient uptake could be an effective strategy to treat cancer. Of course, our data also point towards the need to inhibit multiple pathways to achieve effective cancer therapy and circumvent the metabolic flexibility described here. We hypothesize that in the double mutant tumors deleted for LDHA and GLS, HFSCs failed to upregulate alternative pathways to substitute for the loss of glutaminolysis and lactate utilization as compensatory mechanisms, and as a result, fail to fuel the TCA cycle leading to tumor growth.

## Materials and Methods

### Mice

All animal experiments and related procedures were performed and maintained in accordance with protocols set forth and approved by University of California, Los Angeles (UCLA) Animal Resource Committee (ARC) and the Institutional Animal Care and Use Committee (IACUC) at UCLA in facilities run by the UCLA Department of Laboratory Animal Medicine (DLAM). Animal strains came from Jackson Labs (K15-CrePR, GLS1 fl/fl, LDHA fl/fl, MPC1 fl/fl).

### Two-stage tumorigenesis in mouse skin

Tumors were induced on genetically engineered mice using K15-CrePR animals floxed for either GLS or GLS and LDHA by a cutaneous two-stage skin chemical carcinogenesis[24]. Transgenic animals were shaved and treated with mifepristone (200uL of 10 mg/mL dissolved in filtered sunflower seed oil) daily for 3 days by intraperitoneal injection to delete GLS or GLS and LDHA. After 1 week of treatment with mifepristone, mice were topically treated with a tumor initiating agent, 7,12-Dimethylbenz[*a*]anthracene (DMBA) (400nmol dissolved in acetone). After 1 week of DMBA application, mice were topically treated with a tumor-growth promoting agent, 12-O-tetradecanoylphorbol 13-acetate (TPA), (20nmol dissolved in 100% ethanol) twice a week 3-4 days apart until time of harvest, 25-30 weeks post initial DMBA treatment. Papilloma began to form 10-20 weeks post DMBA treatment and SCCs began to form 20-35 weeks post DMBA treatment.

### Histology, immunofluorescence, and immunohistochemistry

Tumors were harvested from dorsal skin for each indicated genotype and embedded in unfixed OCT compound. Tumors in OCT were cut at 10μm on a Leica 3200 Cryostat for immunostaining, in situ enzymatic activity assays, and hematoxylin and eosin staining. For immunofluorescence staining, slides were briefly fixed in 10% buffered formalin and washed in PBS twice for 10 min. Slides were blocked with 10% goat serum/0.25% Triton-X for 1 hour at room temperature while rotating. The following primary antibodies were used: GLS, KERATIN5, CD45, CD11b, MCT1, MCT4, GLUT1, TXNIP, ASCT2, and p-EGFR. Primary antibodies were diluted into blocking buffer, added to samples, and incubated overnight. The next day, slides were washed in PBS/Tween. Secondary antibodies were added at 1:500 dilution and were incubated on slides rotating at room temperature for 1 hour. Slides were then washed in PBS/Tween, mounted with Prolong Gold with DAPI (Invitrogen), and sealed with clear nail polish. Images were collected at 20x unless otherwise specified. Immunohistochemistry was performed on formalin fixed paraffin-embedded tissue section. Slides underwent antigen retrieval with citrate, were incubated in hydrogen peroxide (30 minutes at 4°C), blocked with 10% goat serum/0.25% Triton-X (1 hour at room temperature) and incubated with primary antibodies (overnight). For detection, we used a secondary horseradish peroxidase-labeled polymer (Dako) and AEC Substrate Chromogen (Vector Laboratories).

| Antibody | Source | Identifier |
| --- | --- | --- |
| anti-rabbit KGA/GAC (GLS) | Proteintech | Cat #: 12855-1-AP |
| anti-chicken KERATIN5 | Biolegend | Cat #: 905901 |
| anti-rabbit CD45 | abcam | ab10558 |
| anti-rat CD11b [M1/70] | abcam | ab197701 |
| anti-rabbit MCT1 | Proteintech | Cat #: 20139-1-AP |
| anti-rabbit MCT4 | Proteintech | Cat #: 22787-1-AP |
| anti-rabbit GLUT1 [EPR3915] | abcam | ab115730 |
| anti-rabbit TXNIP | Invitrogen | Cat #: yf3950376A |
| anti-rabbit ASCT2 (V501) | Cell Signaling | Cat #: 5345 |
| Anti-rabbit EGFR (phosphor Y1068)<br>[EP774Y] | abcam | ab40815 |

### Glutaminase activity assay

Activity of GLS was measure by using GLS activity fluorometric assay kit (Biovision, K455) according to manufacturer’s instructions.

### FDG-PET imaging and analysis

Small-animal PET/CT scans were performed and analyzed as we described in Flores, et al. 2019. SUV was calculated using %ID/g. %ID/g = (SUV divided by animal weight in grams)*100.

### Tracing with ^13^C_6_-D-Glucose, ^13^C_5_-L-Glutamine, and ^13^C_3_-Sodium-L-Lactate

Prior to euthanasia, mice were intraperitoneally infused with ^13^C_6_-D-Glucose (Cambridge Isotope Laboratories, PR-31904), ^13^C_5_-L-Glutamine (Cambridge Isotope Laboratories, PR-30230), and ^13^C_3_-Sodium-L-Lactate (Cambridge Isotope Laboratories, PR-31355) for 10 min. ^13^C_6_-D-Glucose was infused at 2g/kg; ^13^C_5_-L-Glutamine at 0.3mg/g; and ^13^C_3_-Sodium-L-Lactate at 200uL of solution. After 10 min of tracing, tissues were dissected within 3-5 min for metabolite extraction.

### Metabolite Extraction and Liquid Chromatography-Mass Spectrometry

These experiments were performed as previously described in Flores et al. 2019. Briefly, <8mg of fresh tumors were momentarily rinsed in cold 150mM ammonium acetate (pH 7.3) and then added into 1mL of a cold solution of 80% methanol with 10nM trifluoromethanosulfanate. Tumors samples were homogenized with a tissue homogenizer (BeadBug6 model: D1036, 5 cycles, 4000 speed, 30 time) for full homogenization. After removing insoluble material by centrifugation at 17,000g at 4 °C for 10 min, the supernatant was added into a glass vial, and metabolites were dried down under vacuum or EZ-2Elite evaporator. Mass Spectrometry was performed as previously described in Flores et al. 2019. Cell pellets were resuspended in RIPA buffer (Pierce) with Halt protease and phosphatase inhibitors (Thermo-Fisher) on ice. After removing insoluble material by centrifugation at 8000g at 4 °C for 5 min, total protein concentration was determined using the BCA assay kit (Pierce) per manufacturer’s protocol with a microplate reader.

### Statistics and reproducibility / Statistical analysis

All animals used come from a mixed C57BL6/FVB background with no preference in mouse gender for any studies. There was no statistical measure used beforehand to determine sample size. Data were analyzed in Microsoft Excel and GraphPad Prism, and error bars represent SD between two groups performed by a two-tailed t-test analysis. Statistical significances were considered if *pLJ<LJ0.05; **pLJ<LJ0.01; ***pLJ<LJ0.001. Sample size and statistical details can be found in the figure legends.

## Author Contributions

CG, AF, VC, IA, CM, WZ, TT carried out research; CG, AF, HC, and WEL designed experiments and interpreted results; CG and WEL wrote the manuscript; WEL and HC secured resources to carry out the research.

## Acknowledgements

We would like to acknowledge the contributions of those that made this work possible, particularly the staff of core facilities at UCLA including the Genomics Core in the Department of Pathology, the Flow Cytometry Core in the Jonsson Comprehensive Cancer Center, and the Department of Animal Laboratory Medicine (DLAM). We would also like to thank members of the lab and David Shackelford for their thoughtful comments on the manuscript. Schematic figures were created with Biorender.com. The project described was supported by Award Number T32AR071307 from the National Institute of Arthritis and Musculoskeletal and Skin Diseases. The content is solely the responsibility of the authors and does not necessarily represent the official views of the National Institute of Arthritis and Musculoskeletal and Skin Diseases or the National Institutes of Health. This work was also supported by the UCLA Molecular Biology Interdepartmental Doctoral Program, the UCLA Eli and Edythe Broad Center of Regenerative Medicine, the Broad Stem Cell Research Training Program, and the Stem Cell Research Rose Hills Foundation Graduate Scholarship.

## Data Availability

RNA-seq and Metabolomics data will be made available in appropriate database NIH-GEO upon reviewer request, and publicly upon acceptance of the manuscript.

---

**FIGURE S1**

a) RNA-seq data of *WT* (n=5) or *LDHA^KO^* (n=5) tumors showing transcription levels of genes related to glutaminolysis.

**FIGURE S2**

a) *WT* or *GLS^KO^* SCC immunostaining for GLS and KERATIN5, an epidermal marker. Magnified images of strong GLS positive cells in mesenchyme of tumor.

b) *WT* or *GLS^KO^* SCC immunostaining for immune cell surface marker CD45. Magnified images of strong CD45 positive cells in mesenchyme of tumor. Cell nuclei were stained with DAPI.

c) *WT* or *GLS^KO^* SCC immunostaining for macrophage marker CD11b. Magnified images of strong CD11b positive cells in mesenchyme of tumor. Cell nuclei were stained with DAPI.

