## Supplementary figures and images for "Defining metabolic flexibility in hair follicle stem cell induced squamous cell carcinoma"

### Figure S1

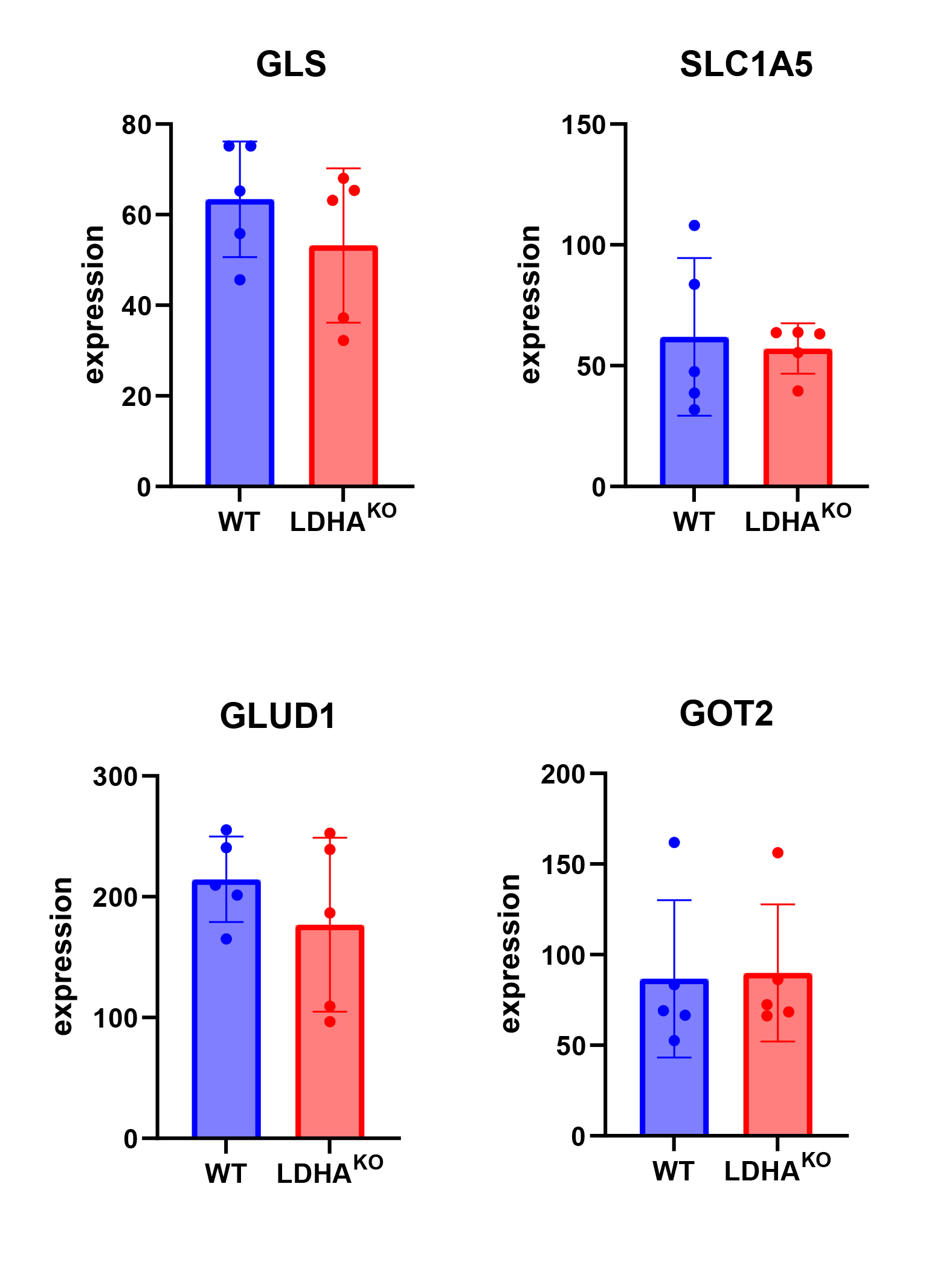

### Figure S2

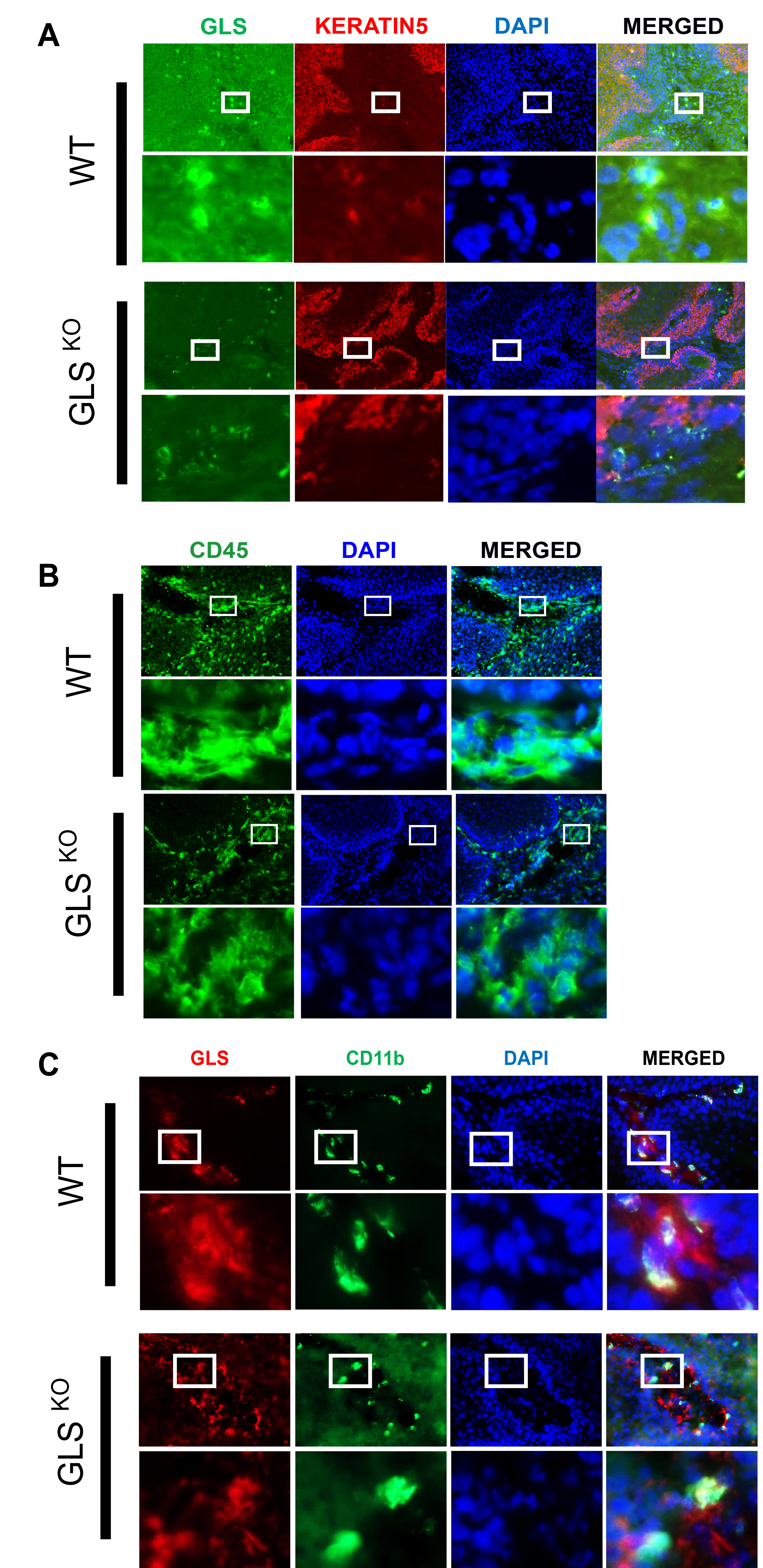
